# Tractostorm: Rater reproducibility assessment in tractography dissection of the pyramidal tract

**DOI:** 10.1101/623892

**Authors:** Francois Rheault, Alessandro De Benedictis, Alessandro Daducci, Chiara Maffei, Chantal M.W Tax, David Romascano, Eduardo Caverzasi, Felix C. Morency, Francesco Corrivetti, Franco Pestilli, Gabriel Girard, Guillaume Theaud, Ilyess Zemmoura, Janice Hau, Kelly Glavin, Kesshi M. Jordan, Kristofer Pomiecko, Maxime Chamberland, Muhamed Barakovic, Nil Goyette, Philippe Poulin, Quentin Chenot, Sandip S. Panesar, Silvio Sarubbo, Laurent Petit, Maxime Descoteaux

**Affiliations:** Sherbrooke Connectivity Imaging Laboratory (SCIL), Université de Sherbrooke, Sherbrooke, Canada; Neurosurgery Unit, Department of Neuroscience and Neurorehabilitation, Bambino Gesù Children’s Hospital, IRCCS, Rome, Italy; Computer Science Department, University of Verona, Verona, Italy; Athinoula A. Martinos Center for Biomedical Imaging, Massachusetts General Hospital and Harvard Medical School, Boston, USA; Cardiff University Brain Research Imaging Centre (CUBRIC), School of Psychology, Cardiff University, Cardiff, United Kingdom; Signal Processing Lab (LTS5), Ecole Polytechnique Fédérale de Lausanne, Lausanne, Switzerland; Department of Neurology, University of California, San Francisco, USA; Imeka, Sherbrooke, Canada; Départment de neurochirurgie, Hôpital Lariboisiere, Paris, France; Department of Psychological and Brain Sciences, Indiana University, Bloomington, USA; UMR 1253, iBrain, Universite de Tours, Inserm, Tours, France; Brain Development Imaging Laboratories, Department of Psychology, San Diego State University, San Diego, California, USA; Learning Research & Development Center (LRDC), University of Pittsburgh, Pittsburgh, USA; ISAE-SUPAERO, Toulouse, France; Department of Neurosurgery, Stanford University, Standford, USA; Division of Neurosurgery, Emergency Department, “S. Chiara” Hospital, Azienda Provinciale per i Servizi Sanitari (APSS), Trento, Italy; Groupe dImagerie Neurofonctionnelle, Institut des Maladies Neurodégénératives – UMR 5293, CNRS, CEA University of Bordeaux, Bordeaux, France

**Author notes:** 2500, boul. de l’Universite, Sherbrooke (Quebec) Canada, J1K 2R1, Email address (Maxime Descoteaux, PhD).

**Keywords:** Diffusion MRI, White Matter, Tractography, Bundle segmentation, Intra-rater, inter-rater, Reproducibility

## Abstract

Investigative studies of white matter (WM) brain structures using diffusion MRI (dMRI) tractography frequently require manual WM bundle segmentation, often called “*virtual dissection*”. Human errors and personal decisions make these manual segmentations hard to reproduce, which have not yet been quantified by the dMRI community. The contribution of this study is to provide the first large-scale, international, multi-center variability assessment of the “*virtual dissection*” of the pyramidal tract (PyT). Eleven (11) experts and thirteen (13) non-experts in neuroanatomy and “*virtual dissection*” were asked to perform 30 PyT segmentation and their results were compared using various voxel-wise and streamline-wise measures. Overall the voxel representation is always more reproducible than streamlines (≈70% and ≈35% overlap respectively) and distances between segmentations are also lower for voxel-wise than streamline-wise measures (¾3mm and ¾ûmm respectively). This needs to be seriously considered before using tract-based measures (e.g. bundle volume versus streamline count) for an analysis. We show and argue that future bundle segmentation protocols need to be designed to be more robust to human subjectivity. Coordinated efforts by the diffusion MRI tractography community are needed to quantify and account for reproducibility of WM bundle extraction techniques in this era of open and collaborative science.

## 1. Introduction

DMRI tractography reconstructs streamlines modeling white matter (WM) connec-tivty. The set of all streamlines forms an object often called the *tractogram* [Jeurissen et al., 2017; Catani and De Schotten, 2008]. When specific hypotheses about known pathways, i.e. WM bundles, are investigated, neuroanatomists design “*dissection plans*” that contain anatomical landmarks and instructions to isolate the bundle of interest from this whole brain tractogram [Catani et al., 2002; Catani and De Schotten, 2008; Chenot et al., 2018; Hau et al., 2016]. Bundles can be segmented to study WM morphology, asymmetries, and then can be associated to specific functions [Lee Masson et al., 2017; Groeschel et al., 2014; Masson et al., 2018; Catani et al., 2007] with approaches similar to other brain structures [Lister and Barnes, 2009; Reitz et al., 2009]. Despite having similar anatomical definitions across publications, the absence of common segmentation protocols for tractography leads to differences that are for the most part unknown and unaccounted for. We need to know how variable our measurements are if we want to be able to have robust tract-based statistics in the future.

The need for a gold standard that quantifies human variability is well-known and well-studied in other fields, such as automatic image segmentation, cell counting or in machine learning [Kleesiek et al., 2016; Entis et al., 2012; Boccardi et al., 2011; Piccinini et al., 2014]. For applications such as hippocampi or tumor segmentation, thorough assessments of reproducibility and multiple iterations of manual segmentation protocols already exist [Boccardi et al., 2015; Frisoni et al., 2015]. These protocols were specifically designed to reduce the impact of human variability and help outcome comparison in large-scale clinical trials across multiple centers [Gwet, 2012; Frisoni et al., 2015].

The reproducibility of manual bundle segmentation will always be lower than manual image segmentation. Image segmentation in 3D requires local decision-making, and the decision to include voxels or not is directly done by raters. However, bundle segmentation requires local decisions that possibly impact the whole volume as streamlines reach outside of the scope of decisions made by raters. Since small hand-drawn regions of interest (ROI) or spheres are used to segment bundles, small mistakes can have far-reaching consequences. Even if ROIs are fairly reproducible in a strict protocol, the resulting bundles could be far from reproducible. This local-decision and global-impact conundrum makes the design of reproducible protocols more difficult and can potentially cause low agreement between raters.

### 1.1. Bundle segmentation

Bundle segmentation is the action of isolating streamlines based on neuroanatomical priors, using known regions where certain conditions need to be satisfied. Inclusion and exclusion regions-of-interests (ROIs) are drawn and defined at the voxel-level using coregistered structural images, and are subsequently used to select the streamlines produced by tractography [Catani et al., 2002; Behrens et al., 2007; Ghaziri et al., 2015; Renauld et al., 2016; Rozanski et al., 2017], as seen in the Figure 1. Streamlines can be included or discarded using inclusion ROIs where streamlines are forced to traverse, and exclusion ROIs that cannot be crossed. Known structures such as grey nuclei, gyri or sulci and recognizable signal signatures can be used as landmarks to create a plan to follow for the segmentation [Catani et al., 2002; Catani and De Schotten, 2008; Hau et al., 2016; Chenot et al., 2018]. In this work, the person performing the task of segmentation (i.e drawing the ROIs, following the protocol) will be referred to as *rater*. Manual segmentation can be performed in software such as, but not limited to, DTI studio [Jiang et al., 2006], Trackvis [Wang et al., 2007], exploreDTI [Leemans et al., 2009], MITK Diffusion [Neher et al., 2012], FiberNavigator [Chamberland et al., 2014], or MI-Brain [Rheault et al., (Figure 1).

**Figure 1:**
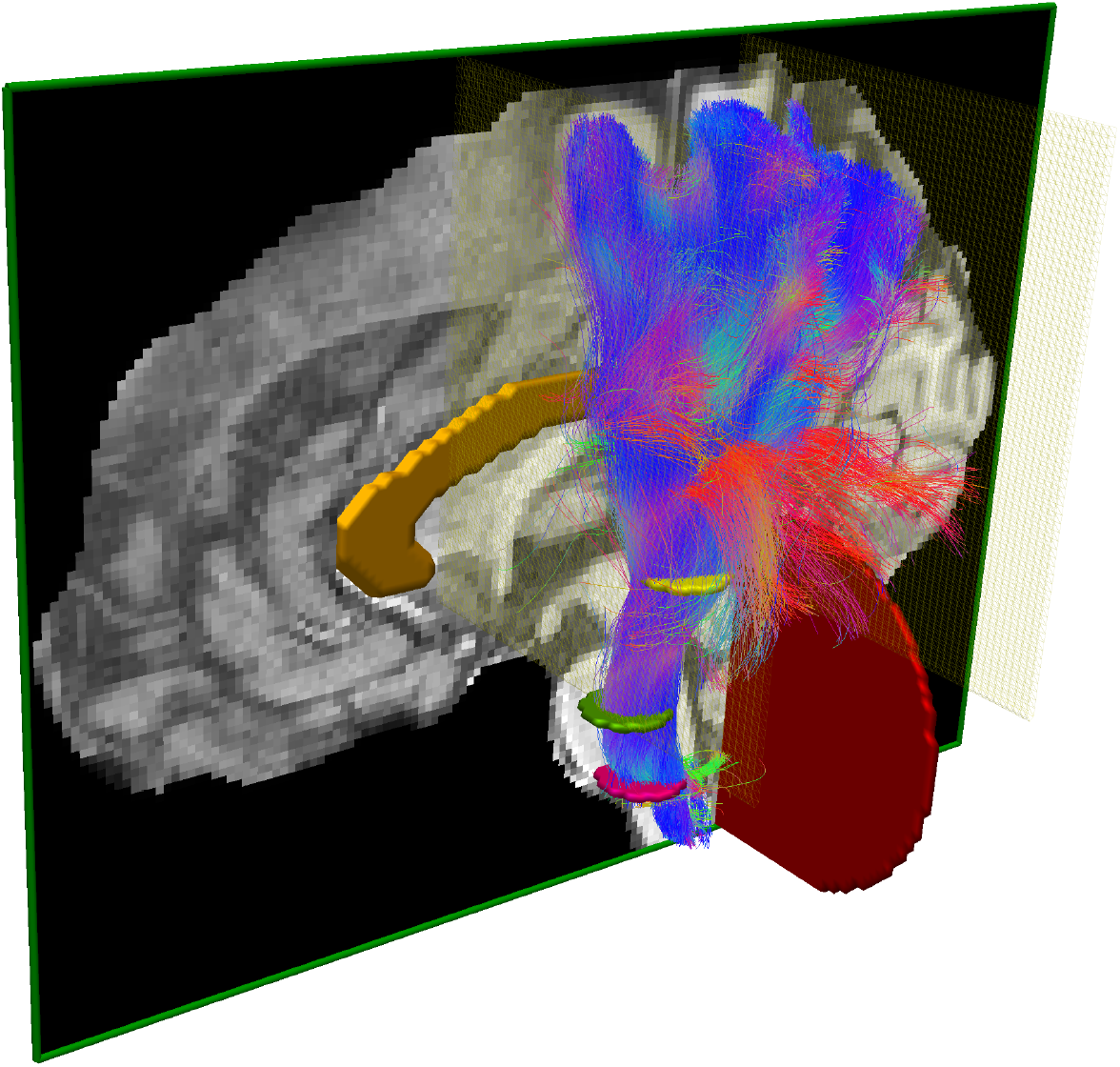
Illustration of the dissection plan of the PyT using the MI-Brain software [Rheault et al., 2016]. 3 axial inclusion ROIs (pink, green, yellow), 1 sagittal exclusion ROI (orange), 2 coronal exclusion ROIs (light yellow) and a cerebellum exclusion ROI (red). The whole brain tractogram was segmented to obtain the left pyramidal tract.

Once a bundle of interest is segmented from a tractogram, the analysis varies according to the research question. It is common to report asymmetry or group difference in bundle volume [Catani et al., 2007; Song et al., 2014; Chenot et al., 2018], diffusion values within the bundle of interest (average fractional anisotropy, mean diffusivity, etc.) [De Erausquin and Alba-Ferrara, 2013; Kimura-Ohba et al., 2016; Ling et al., 2012; Mole et al., 2016] or values along the bundle (called *profilometry* and *tractometry*) [Dayan et al., 2016; Yeatman et al., 2012, 2018; Cousineau et al., 2017]. Spatial distribution of cortical terminations of streamlines can help to identify cortical regions with underlying WM connections affected by a condition [Rushworth et al., 2005; Johansen-Berg et al., 2004; Donahue et al., 2016; Mars et al., 2011; Behrens et al., 2003]. Reporting the number of streamlines (e.g streamline count in connectivity matrix or density maps) is still very much present as a way to compare groups [Jones et al., 2013; Girard et al., 2014; Sotiropoulos and Zalesky, 2017], despite not being directly related to anatomy or connection strength [Jones, 2010; Jones et al., 2013].

### 1.2. Quantifying reproducibility in tractography

When performing segmentation, it is crucial that raters perform the tasks as closely as possible to the dissection plan. Even if a single individual performs all segmentations, the possibility of mistakes or erroneous decisions about landmarks exists [Boccardi et al., 2011; Frisoni et al., 2015; Entis et al., 2012]. High reproducibility is often an assumption, if this assumption is false the consequence could lead to inconsistent outcomes and erroneous conclusions. To assess the level of reproducibility of raters, identical datasets need to be segmented blindly more than once [Gisev et al., 2013; Gwet, 2012; Frisoni et al., 2015]. Reproducibility of segmentations from the same individual is referred to as intra-rater agreement, while reproducibility of segmentation across raters is referred to as inter-rater agreement.

In the field of neuroimaging, voxels are used as the typical representation of data, while the available representation in tractography is in the form of streamlines (i.e. sets of 3D points in space). Figure 2 is a sketch of both representation. Several similarity measures exist to compare voxel-wise segmentations, e.g Dice Score. Most of them have an equivalent formulation to compare sets of streamlines. However, resulting values can widely vary as the spatial distribution is not the same for both representations. Some measures related to streamlines require the datasets to be exactly the same, e.g Dice score, as streamline reconstructions are sets of discrete points with floating point coordinates and not discrete grids like 3D images. For this reason, comparison of streamlines is more challenging and datasets that do not originate from the same source distance in millimeters is often the only available solution [Garyfallidis et al., 2017; Maier-Hein et al., 2017].

**Figure 2:**
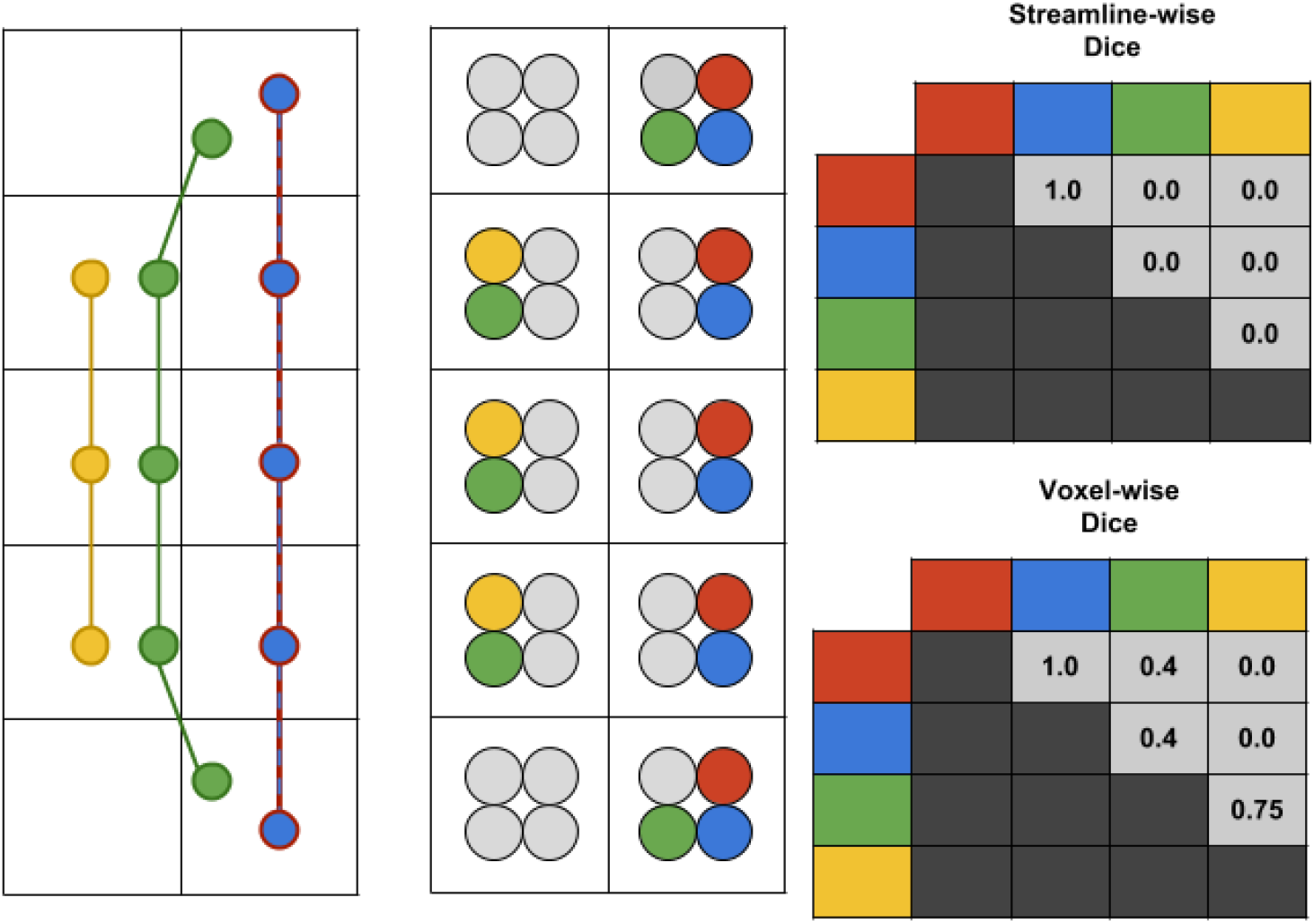
Representation of the Dice Coefficient (overlap) for both the streamline and the voxel representation. For the purpose of a didactic illustration, 4 streamlines are showed in a 2×5 voxel grid, the red and blue streamlines are identical. Each streamline is converted to a binary mask (point-based for simplicity) shown in a compact representation. Voxels with points from 3 different streamlines will results in voxels with 3 different colors, this can be seen as a spatial smoothing. The matrices on the right show values for all pairs (symmetrical). The green and yellow streamline are not identical, which results in a streamline-wise Dice coefficient of zero. However, in the voxel representation they have 3 voxels in common and the result is 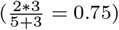.

### 1.3. Summary of contributions of this work

Automatic segmentation methods are becoming more widespread [Guevara et al., 2011; O’donnell et al., 2013; Chekir et al., 2014; Garyfallidis et al., 2017; Zhang et al., 2018; Wasserthal et al., 2018] and aim to simplify the work of raters. The minimal standard of any automatic segmentation method would be to reach the accuracy of raters, thus it is crucial to truly quantify human reproducibility in manual tasks.

The goal of this work is first to quantify human reproducibility of bundle segmentation from dMRI tractography. A measurement of rater (intra and inter) agreement is extremely relevant to set an appropriate threshold for statistical significance. It is also relevant for meta-analysis aiming to study large sets of publications and synthesize their outcomes. An account of human errors or other sources of variability is necessary. A second goal of this work is to investigate overlap, similarity measures and gold standard comparison designed for tractography. Development of easily interpretable measures for bundle comparison is necessary for large datasets. Overall the voxel representation is significantly more reproducible than the streamline representation. The voxel representation is better suited for analysis of tractography datasets (e.g reporting volume instead of streamline count). More details about these different representations and voxel/streamline-wise measures will be detailed in the Method and Results Section.

A thorough approach for bundle comparison quantification gives insights into segmentation quality for future projects. This is needed to facilitate synthesis of findings and outcomes from various publications [Gwet, 2012; Frisoni et al., 2015; Wisse et al., 2017].

## 2. Method

### 2.1. Study design

Twenty-four participants were recruited and divided into two groups: experts and non-experts. The division was based on their neuroanatomical educational background. Participants working as researchers or PhD students in neuroanatomy, neurology or with extended experience in the field performing “*virtual dissection*” as well as neurosurgeons were part of the experts group (11 participants). The non-experts group was composed of MSc, PhD student or Post-Doc in neuroimaging, but without any formal education in neuroanatomy (13 participants). All participants had knowledge of dMRI tractography in general as well as the concept of manual segmentations of tractography datasets. Participation was voluntary and anonymous, recruitment was done individually and participants from various labs in Europe and the USA were solicited. The study was performed according to the guidelines of the Internal Review Board of the Centre Hospitalier Universitaire de Sherbrooke (CHUS).

Five independent tractograms and their associated structural/diffusion images were used, each was triplicated (total of 15). One was untouched, one was flipped in the X axis (left/right) and one was translated. Then, all datasets were randomly named so the tasks could be performed blindly for each participant. Participants were not aware of the presence of duplicated datasets. Five tractotrams were used to obtain stable averages, duplicated datasets were used to score the intra-rater agreement and the multiple participants to evaluate inter-rater agreement. The decision to separate participants in two groups was made to generate additional data about reproducibility in real-life conditions.

Figure 3 shows an overview of the study design. To evaluate intra-rater reproducibility of rater #1, each triplicate was used to compute reproducibility measures. Meaning that 5 (A-B-C-D-E) × 3 (1-2-3) values were averaged to obtain the intra-rater “*reproducibility score*” of a single rater. To evaluate inter-rater reproducibility of rater #1, triplicates were fused and compared to all other raters to obtain a reproducibility measure. Meaning that 5 (A-B-C-D-E) × N (raters) values were averaged to obtain a single rater inter-rater “*reproducibility score*”. To evaluate reproducibility against the gold standard of rater #1 the fused triplicates were also used. Meaning that 5 (A-B-C-D-E) × 1 (gold standard) values were averaged to obtain a single rater gold standard “*reproducibility score*”. The results showed in the Results Section are average values from all raters in each group.

**Figure 3:**
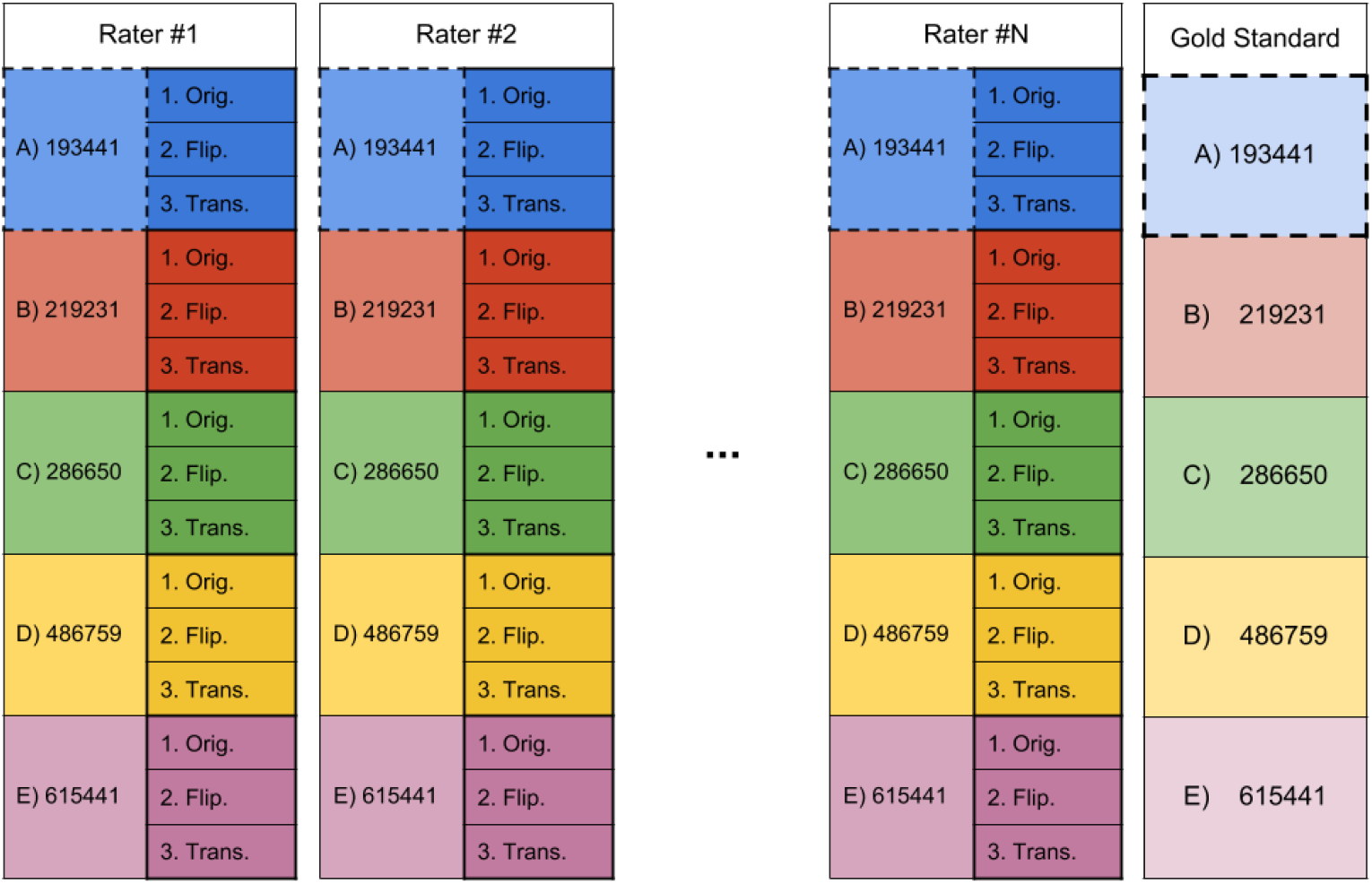
Representation of the study design showing N participants, each received 5 HCP datasets (listed and color-coded) which were replicated 3 times (original, flipped, translated). All participants had to perform the same dissection tasks, on the same anonymized datasets. Intra-rater, inter-rater and gold standard reproducibility were computed using the deanonymized datasets. More details are available in the supplementary materials

### 2.2. DWI datasets, processing and tractography

Tractograms were generated from the preprocessed HCP [Van Essen et al., 2013] DWI data using three shells (1000, 2000, 3000) with 270 directions. The B0, fractional anisotropy (FA) and RGB (colored FA) images were computed from DWI to be used as anatomical reference during segmentation. Constrained spherical deconvolution (CSD) using a FA threshold from a tensor fit on the b=1000s/*mm*^2^ was used to obtain fiber orientation distribution functions (fODF) [Tournier et al., 2007; Descoteaux et al., 2007] (spherical harmonic order 8) from the b=2000s/*mm*^2^ and b=3000s/*mm*^2^ shells. Probabilistic particle filtering tractography [Girard et al., 2014] was subsequently computed at 30 seeds per voxel in the WM mask (FSL FAST [Woolrich et al., 2009]) to make sure sufficient density and spatial coverage were achieved.

The CSD model was also used for bundle-specific tractography (BST) to further improve density and spatial coverage of the bundle of interest [Rheault et al., 2018; Chenot et al., 2018]. This was to ensure that the full extent of the CST was reconstructed and to ensure not to have criticisms from our experts in neuroanatomy complaining of missing CST parts. A large model that approximates the CST was used to generate streamlines with a strong preference for the Z axis (up-down). For BST, the same tractography parameters were used except for seeding, which was exclusively done from the precentral gyrus, postcentral gyrus and brainstem at 5 seeds per voxel.

The whole brain tractogram and the CST-specific tractogram were fused. To accommodate all participants and the wide range of computer performance, tractograms were compressed using a 0.2mm tolerance error [Rheault et al., 2017; Presseau et al., 2015] and commissural streamlines were removed and datasets split into hemispheres.

### 2.3. Dissection plan and instructions

Each participant received their randomly named datasets, a document containing instructions for the segmentation and a general overview of a segmentation as example (see supplementary materials). The segmentation task consisted in 15 segmentations of the pyramidal tract (left and right). Segmentation involved using 3 WM inclusion ROIs (Internal capsule, Midbrain and Medulla Oblongata) and 2 exclusion ROIs (one plane anterior to the precentral gyrus and one plane posterior to the postcentral gyrus). The detailed segmentation plan is available in the supplementary materials [Chenot et al., 2018].

Participants had to perform the segmentation plans, following the instructions as closely as possible. The dataset order was provided in a spreadsheet file. Participants had to choose between two software; Trackvis [Wang et al., 2007] or MI-Brain [Rheault et al., 2016]. This decision was made to guarantee that the data received from all participants was compatible with the analysis.

Metadata such as date, starting time and duration had to be noted in the spreadsheet file. Upon completion, the participants had to send back the same 15 folders with two tractography files in each, the left and right pyramidal tract (PyT).

### 2.4. Bundles analysis

Once returned by all participants, datasets were de-randomized to match triplicates across participants. The duplicates (flipped and translated) were reverted back to their native space and all datasets (images and tractograms) were warped to a common space (MNI152c 2009 nonlinear symmetrical) using the Ants registration library [Fonov et al., 2011; Avants et al., 2008] to simplify the analysis. With all datasets having a uniform naming convention and in a common space, the intra-rater and inter-rater reproducibility can be assessed.

#### Individual measures

Reproducibility can be assessed using various measures. Volume and streamline count are the main attributes obtained directly from files. They do not provide direct insight about reproducibility, but one could expect that very similar segmentations should have very similar values. However, this does not provide any nuance or specific information about difference. In this work results for the left and right PyT are averaged together without distinction, they are considered the same bundle during the analysis.

#### Intra-rater and inter-rater

Each participant performed the same tasks on each triplicate. The goal of this triplication is to evaluate intra-rater reproducibility. Since all participants had access to the same datasets, inter-rater reproducibility can be assessed too.

Computing the average value from all pairwise combinations provides an estimate of the agreement between multiple segmentations of a same bundle. The deviation can also provide insights about the consistency of these segmentations. Measurement values can be between 0 and 1, such as Dice and Jaccard [Dice, 1945], meaning they are independent of the size. An alternative to overlap measures are similarity measures, which can provide insights about the distance between two segmentations (in millimeter). Even when spatial overlap between two segmentations is low, both can be very similar in shape [Descoteaux et al., 2004; Garyfallidis et al., 2010]. Figure 5 shows bundles and how to interpret these measures. Pearson’s correlation coefficient obtained from density maps provides insight into the statistical relationship and spatial agreement between two segmentations [Hyde and Jesmanowicz, 2012]. More details on available measures for tractography are available in the supplementary materials.

**Figure 4:**
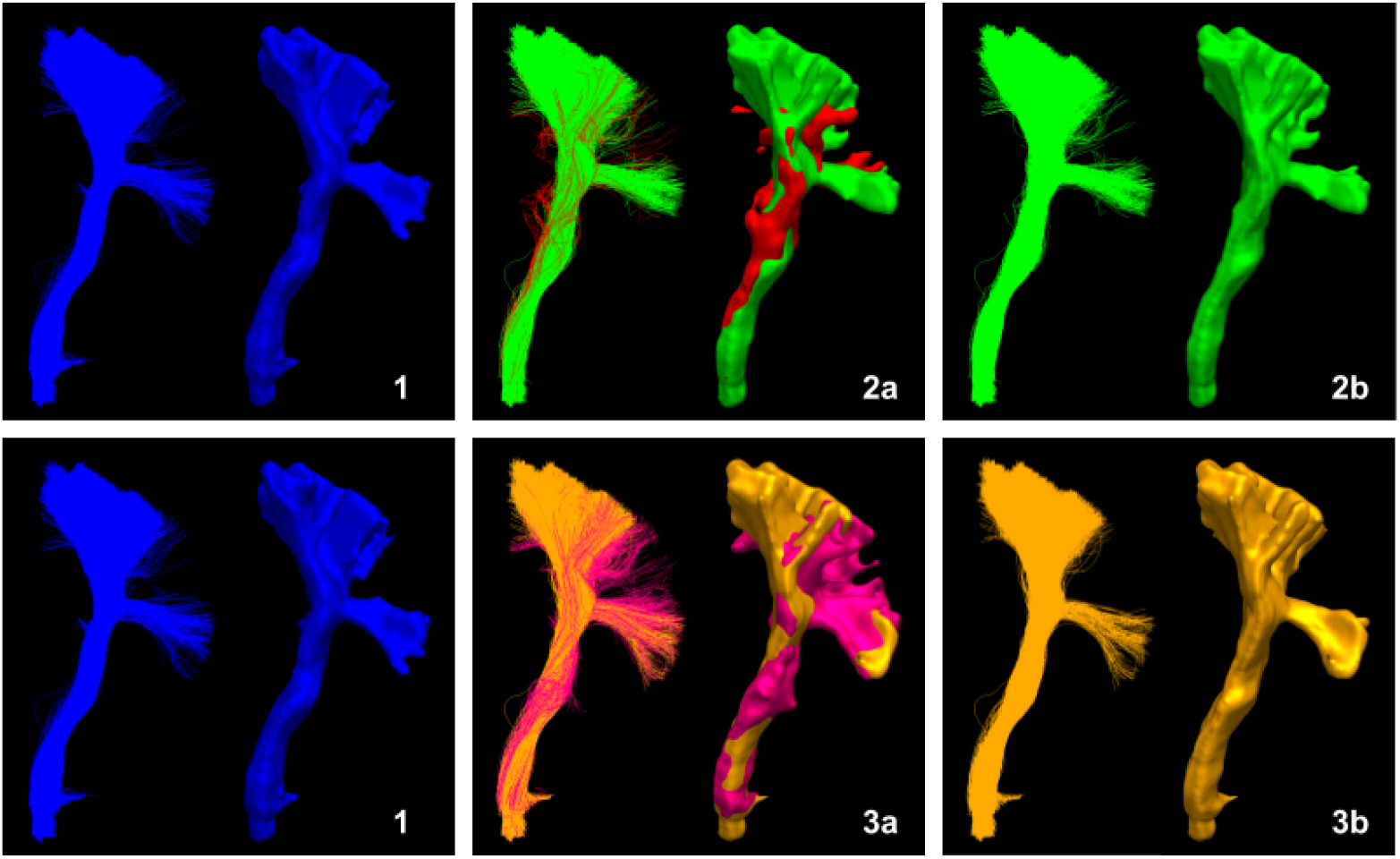
Comparison of bundles and the impacts of spurious streamlines on the reproducibility measurements. Each block shows streamlines on the left and the voxel representation on the right (isosurface). Block 2a and 3a shows the core (green/orange) and spurious (red/pink) portion of the bundle. Block 2b and 3b only shows the core portion of the bundle.

**Figure 5:**
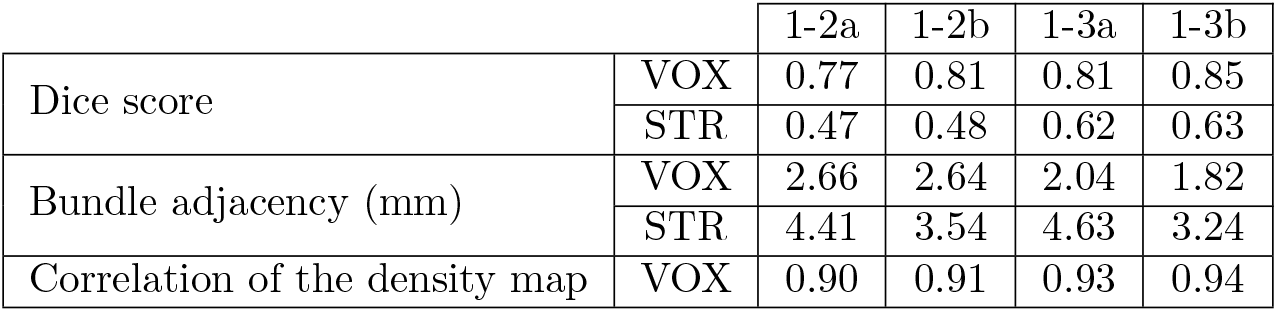
Table showing the reproducibility “*score*” between bundles, VOX marks voxel-wise measures and STR marks streamline-wise measures.

The most insightful measures are represented by the overlap (Dice coefficient), distance (bundle adjacency) and density and spatial coherence (density correlation). Each measure provides a way to interpret the data at hand, but there is no single true measure to summarize intra-rater and inter-rater agreement. Multiple measures were computed and are all available in the supplementary materials along more detailed description for each of them.

#### Gold standard

When multiple raters provide segmentations from an identical dataset, it is of interest to produce a gold standard. For a voxel representation, a probability map can be constructed, where each voxel value represents the number of raters that counted the voxel as part of their segmentation [Frisoni et al., 2015; Iglesias and Sabuncu, 2015; Langerak et al., 2015; Pipitone et al., 2014]. This can be normalized and then thresholded to obtain a binary mask representing whether or not the voxel was segmented by enough rater. A threshold above 0.5 is often referred to as a majority vote. The same logic can be applied to streamlines, each streamline can be assigned a value based on the number of raters that considered it part of their segmentation.

This can be seen in Figure 6 where increasing the minimal vote threshold reduces the number of outliers and overall size. In this work, the gold standard *does not* represent the true anatomy and should not be interpreted as such. It simply represents the average segmentation obtained from a tractogram.

**Figure 6:**
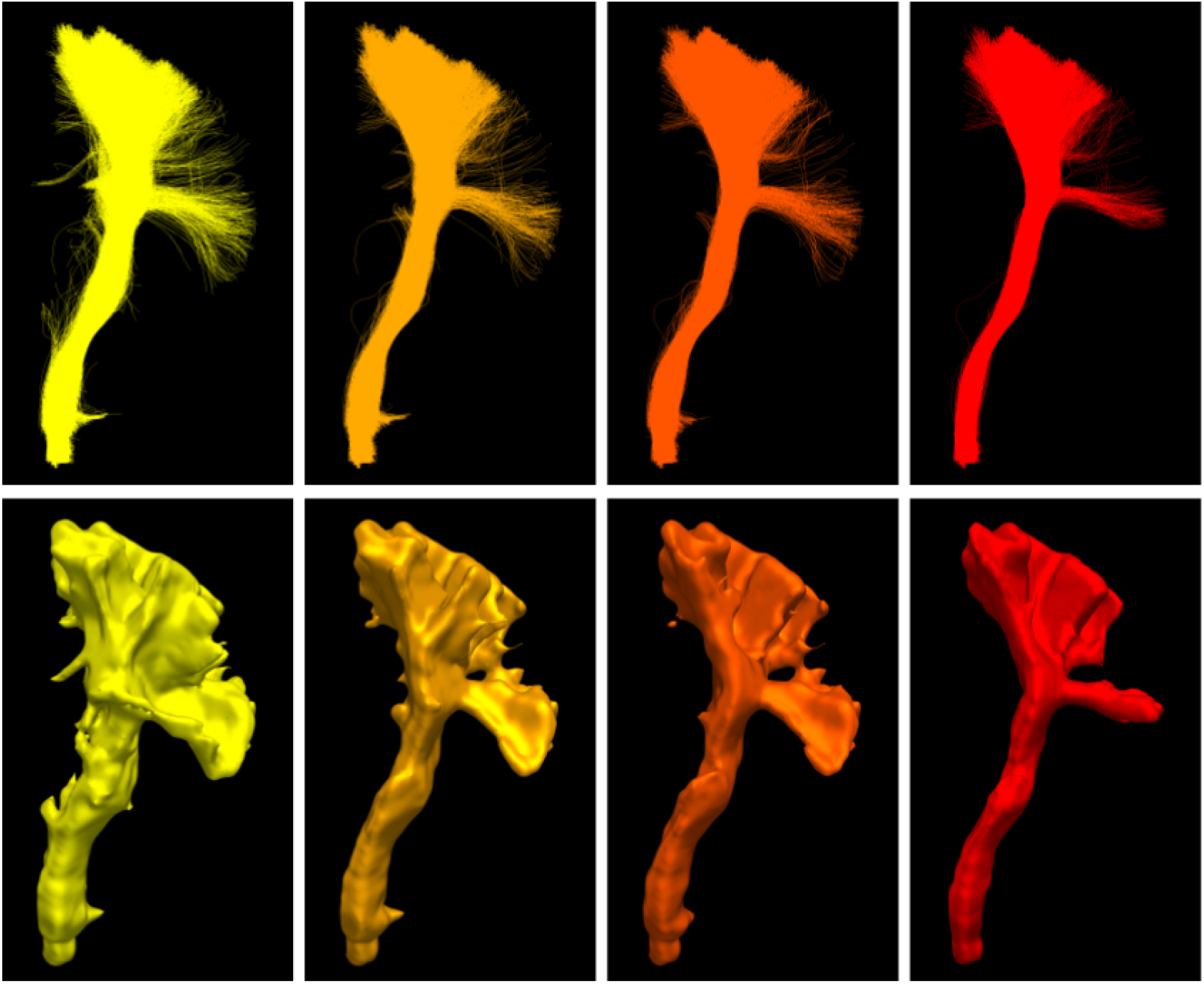
Gold standard obtained from 7 segmentations, first row shows the streamline representation and the second row shows the voxel represented as a smooth isosurface. From left to right, multiple voting ratios were used 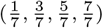, each time reducing the number of streamlines and voxels consider part of the average segmentation. A minimal vote set at 1 out of 7 (left) is equivalent to a union of all segmentations while a vote set at 7 out of 7 (right) is equivalent to an intersection between all segmentations.

All elements that are not in a gold standard are true negatives and all the ones present are true positives. By construction, the gold standard does not contain false positives or false negatives. Binary classification measures are available such as sensitivity or specificity. However, several other measures are available and each are a piece of the puzzle leading to a more accurate interpretation [Garyfallidis et al., 2017; Chang et al., 2009; Schilling et al., 2018].

To produce our gold standard a majority vote approach was used from the segmentations of the experts group, as their knowledge of anatomy was needed to represent an average version of the bundle of interest. The vote was set at 6 out of 11 and each of the 5 datasets got its own left and right gold standard. Since the representation at hand is streamlines (which can be converted to voxels), a streamline-wise and a voxel-wise gold standard were created.

## 3. Results

On average, experts produce “*smaller*” bundles than non-experts, their volume and streamline count is lower than non-experts, as it can be observed in Table 1 and Figure 7. This difference between groups is statically significant (*p – value* < 0.01). In the following sections, all explicit comparisons between groups are statistically significant using a standard Welch’s t-test for the means of two independent samples, which does not assume equal population variance (*p* – *value* < 0.01). The range of values for segmentation measures is wider for non-experts, meaning that either intra-rater or inter-rater variability is higher. As mentioned earlier, this is useful insight about reproducibility, but lacks nuance and context.

**Figure 7:**
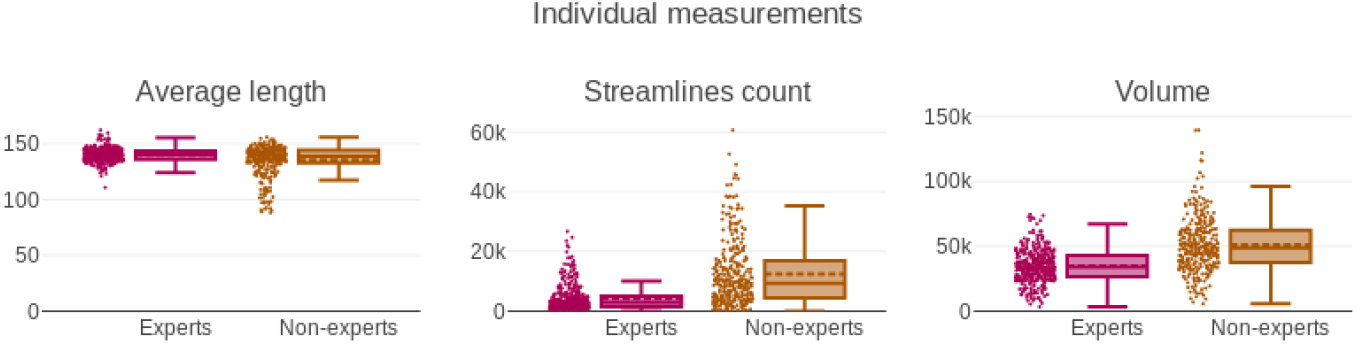
Boxplots and scatter plots showing distribution of the 3 measurements related to individual files for both groups.

**Table 1:** Table showing main values from boxplots of the 3 measurements related to individual files for both groups. The columns show the average value and the standard deviation for each group. VOX marks voxel-wise measures and STR marks streamline-wise measures. Rows shown in bold mean that the two groups (experts and non-experts) do not have the same distribution.

|  |  | Expert |  | Non-experts |  |
| --- | --- | --- | --- | --- | --- |
|  |  | Mean | STD | Mean | STD |
| Volume ($mm^3$) | **VOX** | 34835 | 12625 | 51146 | 20966 |
| Streamline count | **STR** | 4331 | 4457 | 12489 | 11091 |
| Mean length (mm) | **STR** | 140.33 | 7.81 | 138.70 | 11.29 |

### 3.1. Intra-rater evaluation

All reported values can be seen in Table 2 and in Figure 8. The average intra-rater overlap is represented by the voxel-wise Dice coefficient and is on average 0.72 for experts and 0.78 for non-experts. Streamline-wise Dice coefficient is much lower at 0.31 and 0.52 for both groups respectively. A higher Dice score value means that participants of a group are, on average, more reproducible with themselves. Non-overlapping voxels are on average at a distance 2.13mm for experts and 2.58mm for non-experts (lower Mean value represent higher similarity). Streamline-wise distance is lower in the experts group at 5.27mm while the non-experts group is at 6.12mm. The average density correlation is equal for both group at 0.82 and 0.82 for the experts and non-experts group respectively (*p — value* > 0.01).

**Figure 8:**
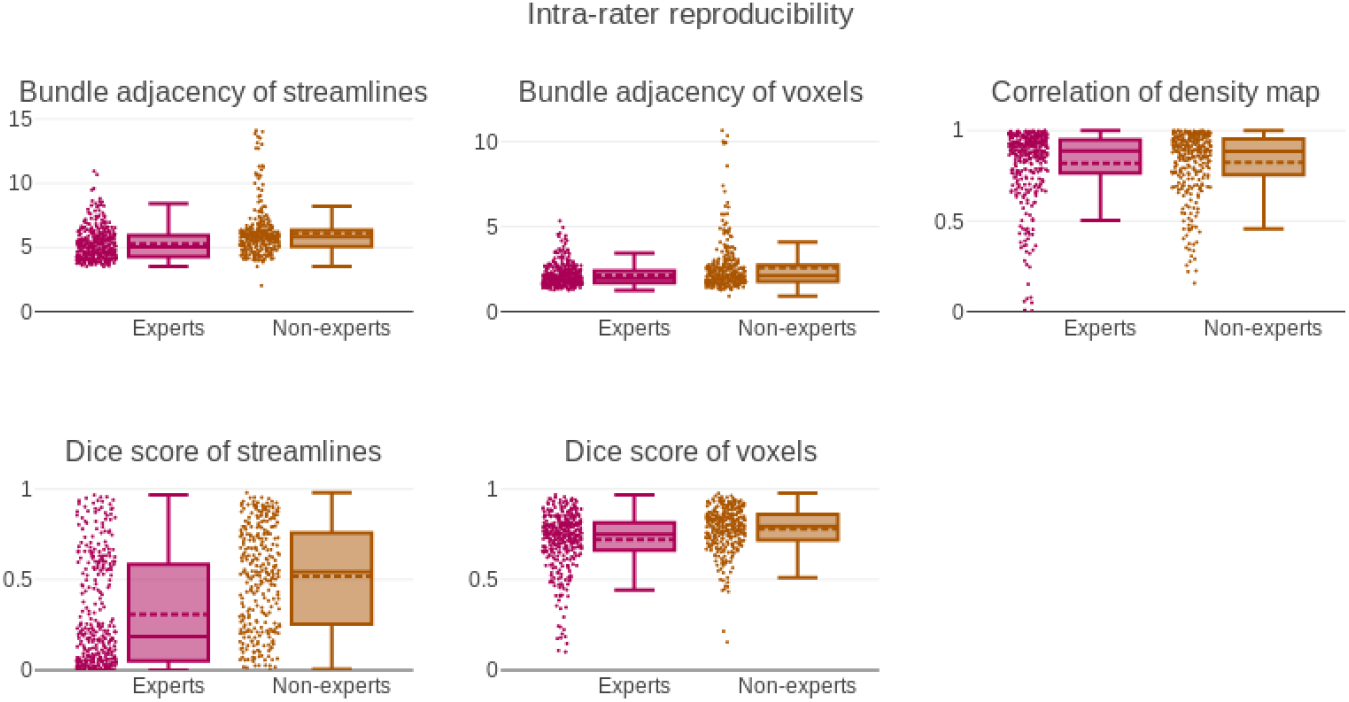
Boxplots and scatter plots showing distribution of the 3 measurements related to pairwise comparison measures for intra-rater segmentations.

**Table 2:** Table showing main values from boxplots of the 3 measurements related to pairwise comparison measures for intra-rater segmentations. Voxel and streamline values of the same measures are in the same cell. Rows shown in bold mean that the two groups (experts and non-experts) do not have the same distribution.

|  |  | Expert |  | Non-experts |  |
| --- | --- | --- | --- | --- | --- |
|  |  | Mean | STD | Mean | STD |
| Dice score | **VOX** | 0.72 | 0.16 | 0.78 | 0.12 |
|  | **STR** | 0.31 | 0.30 | 0.52 | 0.28 |
| Bundle adjacency (mm) | **VOX** | 2.13 | 0.66 | 2.58 | 1.53 |
|  | **STR** | 5.27 | 1.26 | 6.12 | 1.89 |
| Correlation of density map | VOX | 0.82 | 0.20 | 0.82 | 0.18 |

### 3.2. Inter-rater evaluation

To minimize the influence of intra-rater reproducibility during the evaluation of interrater reproducibility, the triplicate datasets were fused into a single bundle. This was performed to approximate the results as if participant segmentations had no intra-rater variability. This lead to a underestimation of inter-rater variability, but necessary to separate source of variability later in the analysis. Voxel-wise Dice coefficient is on average higher between experts than between non-experts, at 0.75 and 0.67 respectively. Streamline-wise Dice coefficient is not statistically different (*p – value >* 0.01) at 0.34 and 0.32. Voxel-wise distance is on average lower for the experts group than non-experts, 2.74mm and 3.85mm respectively. The average density correlation is higher between experts at 0.88 while non-experts are at 0.71. The standard deviation is higher for the non-experts group, meaning that the similarity among non-experts is not only lower on average, but widely varies. All reported values can be seen in Table 3 and in Figure 9.

**Figure 9:**
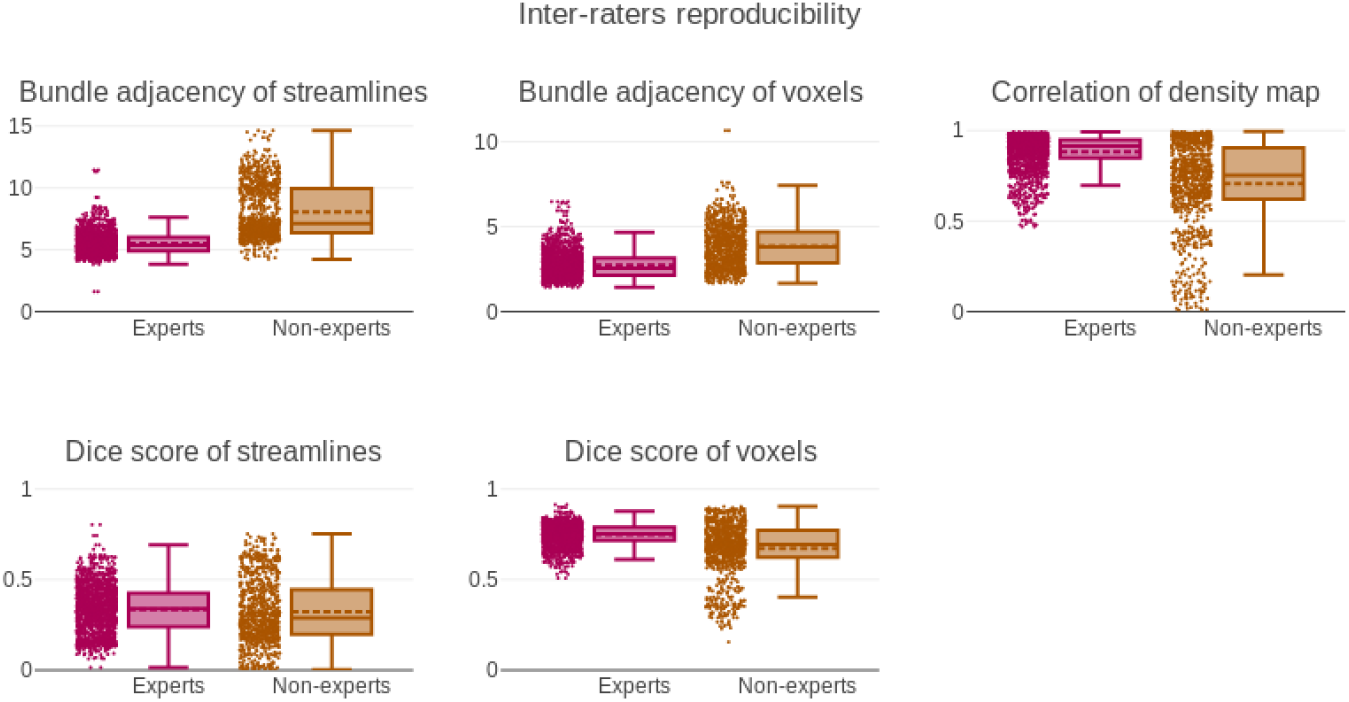
Boxplots and scatter plots showing distribution of the 3 measurements related to pairwise comparison measures for inter-rater segmentations.

**Table 3:**
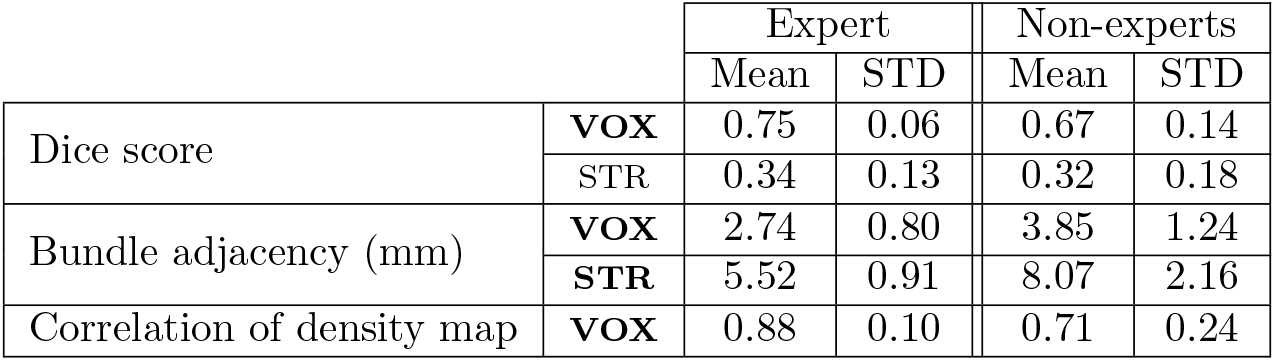
Table showing main values from boxplots of the 3 measurements related to pairwise comparison measures for inter-rater segmentations. Voxel and streamline values of the same measures are in the same cell. Rows shown in bold mean that the two groups (experts and non-experts) do not have the same distribution.

### 3.3. Gold standard evaluation

All reported values can be seen in Table 4, 5 and in Figure 10, 11. Comparisons to the computed gold standard shows that on average experts and non-experts obtain segmentation roughly similar to the average segmentation. However, all measures show that segmentations from experts are on average closer to the gold standard than those of non-experts. This was expected as the gold standard was produced using segmentations from the experts group. Values for streamline-wise measures are lower for Dice coefficient and density correlation and higher for bundle adjacency, meaning that reproducibility is harder to achieve using the streamline representation. This was a similar trend observed in intra-rater and inter-rater values.

**Figure 10:**
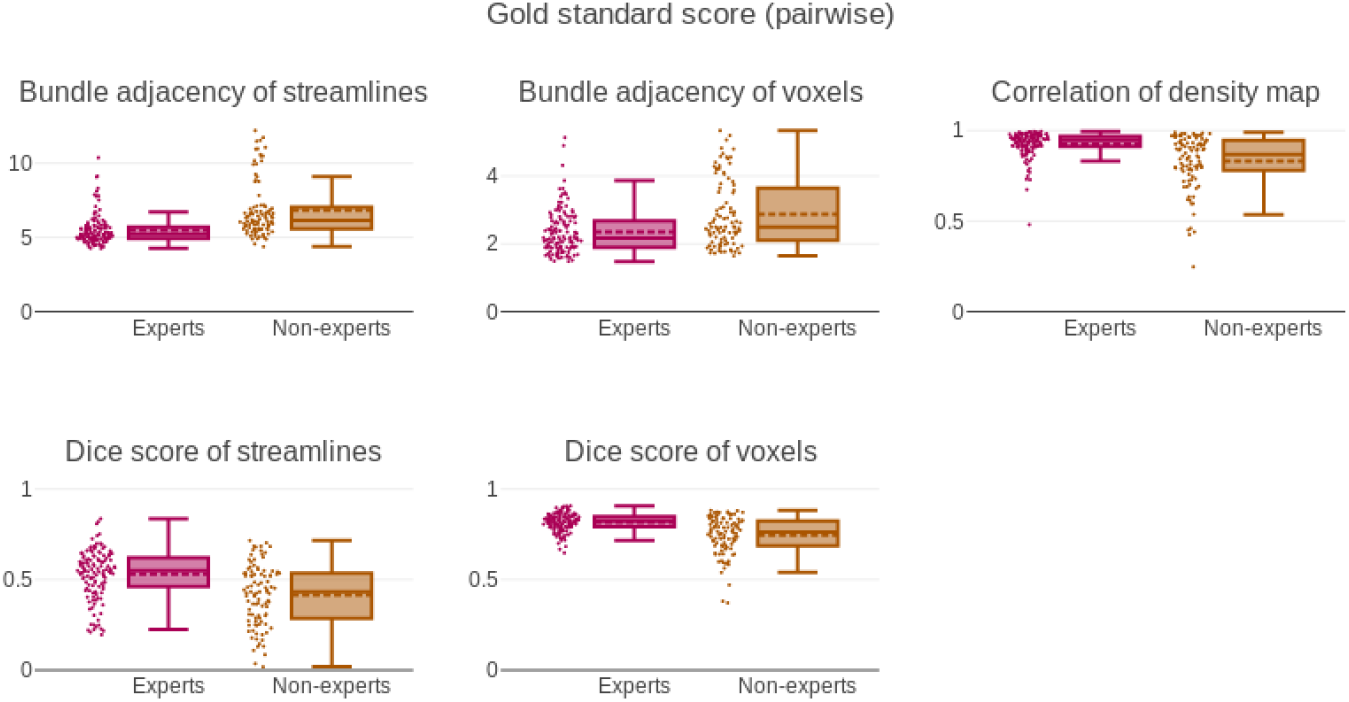
Boxplots and scattera plots showing distribution of the 3 measurements related to pairwise comparison measures against the gold standard.

**Table 4:**
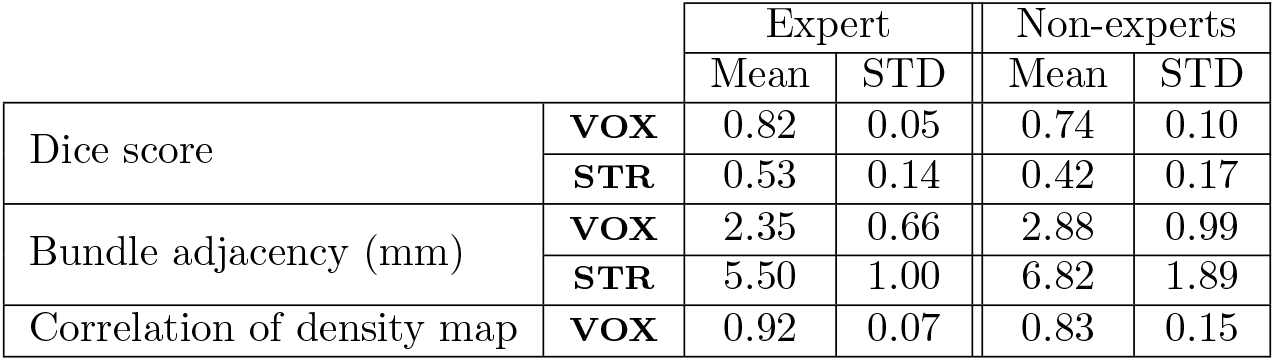
Table showing main values from boxplots of the 3 measurements related to pairwise comparison measures against the gold standard. Voxel and streamline values of the same measures are in the same cell. Rows shown in bold mean that the two groups (experts and non-experts) do not have the same distribution.

|  |  | Expert |  | Non-experts |  |
| --- | --- | --- | --- | --- | --- |
|  |  | Mean | STD | Mean | STD |
| Dice score | **VOX** | 0.82 | 0.05 | 0.74 | 0.10 |
|  | **STR** | 0.53 | 0.14 | 0.42 | 0.17 |
| Bundle adjacency (mm) | **VOX** | 2.35 | 0.66 | 2.88 | 0.99 |
|  | **STR** | 5.50 | 1.00 | 6.82 | 1.89 |
| Correlation of density map | **VOX** | 0.92 | 0.07 | 0.83 | 0.15 |

**Table 5:**
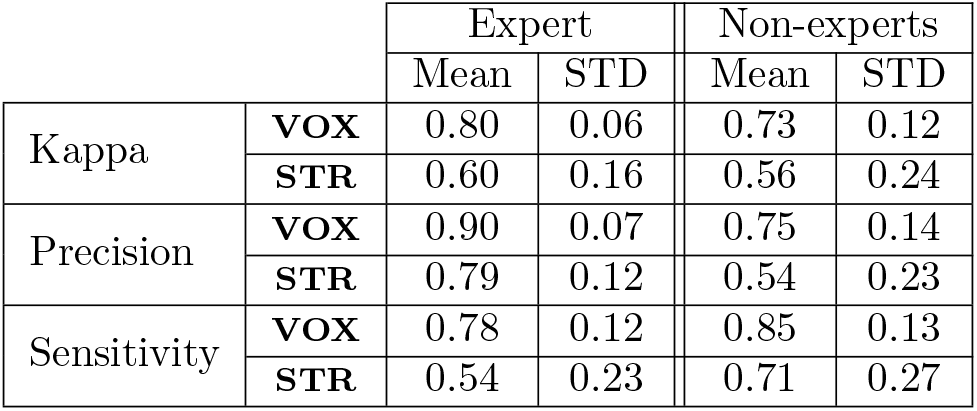
Table showing main values from boxplots of the 3 measurements related to binary classification measures against the gold standard. Rows shown in bold mean that the two groups (experts and nonexperts) do not have the same distribution.

Specificity and accuracy reach above the 95% for both groups both for streamlines or voxels. Meaning that experts and non-experts alike classified the vast majority of true negatives correctly. Since specificity is near a value of 1.0, the Youden score is almost equal to sensitivity. All 3 measures take into account the true negatives, which far outweigh the true positives, in our datasets, for this reason they were removed from Figure 11 and shown only in the supplementary materials. Sensitivity is much lower at 0.59 and 0.71 for experts and non-experts respectively, as both groups partially capture the gold standard. Precision is higher for experts than for non-experts, meaning that experts were providing segmentations approximately the same *size* as the gold standard while non-experts were providing much bigger segmentations (that generally encompass the gold standard). This explains the higher sensitivity and lower specificity of nonexperts. The average Kappa and Dice score is lower for experts at 0.67 and 0.72 while the non-experts average is 0.69 and 0.73, respectively. The Kappa score takes into account overlap with the probability of randomly obtaining the right segmentation. Given the dimensionality of our data, getting the right segmentation by accident is very low, explaining why the Kappa and Dice score are very similar. It is important to consider that the ratio of true negatives to true positives is not the same for both representations (voxels vs. streamlines).

**Figure 11:**
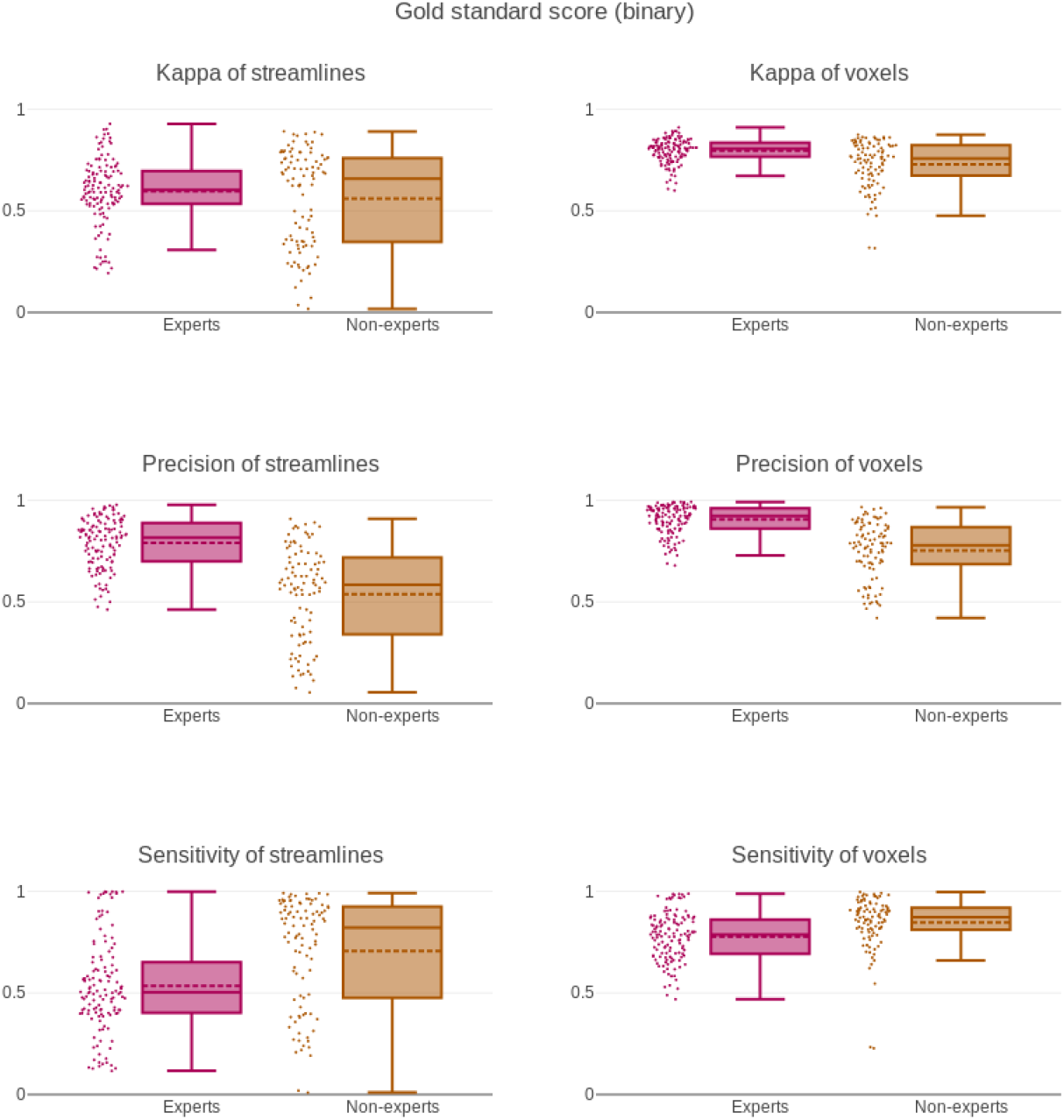
Boxplots and scatter plots showing distribution of the 6 measurements related to binary classification measures against the gold standard.

The computation of inter-rater reproducibility was performed using the fused triplicate to minimize the influence of intra-rater reproducibility. The approach to fuse the triplicate is simply an approximation, producing more than 3 segmentations of the same datasets would be necessary to perfectly evaluate intra-rater reproducibility. However, the 5 datasets used for this study represent sufficiently similar tasks to consider our approximation adequate for this work. Preliminary analysis showed low correlation values, between a participant “*score*” for intra-rater reproducibility and inter-rater reproducibility. Correlation was between 0.2 and 0.4 for all measures, this indicates that there is no clear link between the reproducibility of a participant’s own segmentations and the agreement with other participants.

## 4. Discussion

### 4.1. Evaluation of protocols

This work illustrates that intra-rater and inter-rater agreement is far from perfect even when following a strict and “*simple*” segmentation protocol. The intra-rater and inter-rater agreement must be taken into account when researchers compare bundles obtained from manual segmentations. When human expertise is required for a project, it is crucial that people involved in the manual segmentation process evaluate their own reproducibility, even if they have sufficient neuroanatomy knowledge and extensive experience in manual segmentation. This measure of error will likely increase the threshold for statistical significance. In such case, either more datasets will be needed, or a better protocol for segmentation needs to be designed [Gwet, 2012; Boccardi et al., 2015]. The similarity between both groups indicates that with the right protocol, it would be possible to train people without anatomical background to perform tasks with results and quality similar to experts.

Without such evaluation it is impossible for experts and non-experts alike to know how reproducible they are beforehand. Since their “*scores*” vary with the protocol, the bundle of interest and possibly other factors, it is important to consider an evaluation before performing large-scale segmentation procedure [Frisoni et al., 2015]. An alternative to guarantee more reproducible results is to design an appropriate protocol for nonexperts and to perform tasks blindly more than once. The results can then be averaged, which will make outliers and errors easier to be identified.

This study did not allow for collaboration and did not micro-manage participants, meaning they were left with the instructions without further intervention from the organisers. In a situation where a segmentation plan can be defined in groups and techniques can be improved along iterations of the plan, the intra-rater and inter-rater agreement would likely go up. This study aimed at the evaluation of participants following instructions from a protocol, similar to the ones present in books, publications or online examples.

### 4.2. Measures and representations

In this work the intra-rater agreement was higher for non-experts than experts, without more information we could have concluded that non-experts were more meticulous when they were performing their manual segmentations. However, by looking at sensitivity and precision we can see than non-experts had “*bigger*” segmentations. Experts are likely stricter in their decision-making process, this could amplify the local-decision and global-impact conundrum mentioned earlier. A more liberal, less rigid, segmentation likely makes it easier to be reproducible, but does not necessarily make it valid. This is an example showing the importance of having more than one type of measure to obtain a complete picture.

In tractography, it is common to use a single measure to portray a complex phenomenon. Most measures used are simplified to have easily interpretable results. The previous example shows the importance of using more than one type of measurements to obtain a complete picture of the reproducibility. Reproducibility “*scores*” are likely to vary with the project and the bundle of interest. This needs to be addressed as a community. The discrepancy between protocol quality, reproducibility and conclusion put forward in the literature can be problematic.

For binary measures (accuracy and specificity), scores were both above 95% as it is easy to discard true negatives, and consequently did not provide much insight. Similarly to the curse of dimensionality in machine learning [Verleysen and Francois, 2005; Ceotto et al., 2011], our datasets typically contain millions of voxels (or streamlines), of which only a few thousands true positives are considered during segmentation. Thus, the vast majority of true negatives are rapidly discarded resulting in both accuracy and specificity almost reaching 100%. Sensitivity provides more information, as true positives are more difficult to get, since they are rarer in the tractograms (few thousands out of millions) [Maier-Hein et al., 2017]. This needs to be taken into account using precision, as in some cases, strict segmentation is encouraged because false positives are more problematic than false negatives. Streamline-wise measures show lower agreement, meaning that reproducible results are likely more difficult to achieve with the streamlines representation.

More complex measures need to be designed to fully capture the complexity of tractography datasets and compare them, even across datasets or for longitudinal studies. Currently, more advanced measures that capture fanning, spatial coherence, localized curvature and torsion or spectral analysis are still rare, despite being used in other neuroimaging fields [Esmaeil-Zadeh et al., 2010; Lombaert et al., 2012; Glozman et al., 2018; Cheng and Basser, 2018].

### 4.3. Tractography algorithms

Iterative tracotography algorithms are commonly divided in two categories: Deterministic or probabilistic [Tournier et al., 2012; Garyfallidis et al., 2014]. The most striking difference between both approaches is that probabilistic pathways cover more volume, as they can easily curve and explore more ground. On the other hand, deterministic will be more conservative due to curvature restrictions, thus leading to less exploration and therefore smaller volume [Maier-Hein et al., 2017].

Manual segmentation of deterministic tractograms is likely more reproducible, since small differences in ROI placement will have a smaller impact on the resulting bundle. The local-decision and global-impact conundrum mentioned earlier is more obvious with probabilistic tractography. Other tractography algorithms, such as global tractography [Kreher et al., 2008; Mangin et al., 2013; Christiaens et al., 2015; Neher et al., 2012], are likely to have different reproducibility “*scores*”, even with the same segmentation protocol. Any change to the preprocessing could lead to unexpected change in the reproducibility “*scores*”. Using the same datasets and tractography algorithm, but increasing or decreasing the number of streamlines could also change the reproducibility “*scores*”. Investigations of other bundles of interest would likely lead to different reproducibility “*scores*”, using another anatomical definition of the PyT or even having the anatomical definition taught to participants would have the same effect. However, the general conclusion remains that reproducibility needs to be quantified for specific projects and protocols. Reproducibility “*scores*” cannot be generalized and any attempt would be futile.

### 4.4. Impact on analysis

If variability needs to be minimized further than the defined protocol, a simple recommendation is to have a single rater perform each task multiple times or multiple raters perform each task multiple times (or a subset of tasks). This way, it is guaranteed that each dataset is segmented more than once, decreasing the error risk. Regardless of the decision made, it is of great importance to quantify the reproducibility of manual segmentation of raters involved in the project before doing any statistics or group comparisons. This could drastically change the statistical significance of results. As of now, to the best of our knowledge, human variability and errors are rarely considered. Sources of variability needs to be accounted to truly enable synthesis of work across multiple centers. Even when automatic or semi-automatic methods are used, they first need to be evaluated with agreed upon measures and reach or surpass human standards.

The extension to other bundles of interest or other segmentation plans is not trivial and the only conclusion that stands is that agreement is never 100% and that a unique measure is not enough to represent the whole picture for tractography segmentation. The desire to simplify measures or have only one value to describe quality or reproducibility of segmentations needs to be discouraged. The nature of our datasets makes this task much more complex to interpret than 2D or 3D images, and it is imperative that the field comes to understand and agree on measures to report. This is more relevant than ever as the field grows and now that open science is becoming more popular and reproducibility studies are encouraged. Similarly to other neuroimaging fields, such as hippocampi segmentation, standardized protocols need to be developed and designed to be used across multiple centers without active collaboration or micromanagement.

### 4.5. Future work

Future work includes the creation of a database containing bundle segmentations and metadata from participants that will be available online so further analysis can be done. As for now, a preliminary upload of the participants segmentation is available on Zenodo (https://doi.org/10.5281/zenodo.2547025), which will be updated. In this work, metadata was not used to evaluate duration as a variable influencing reproducibility. Investigating the relationship between variability and duration of a task or looking for bias (inter-hemispheric or software influence). An online platform similar to the Tractometer [Côté et al., 2013] or a Nextflow pipeline [Di Tommaso et al., 2017] is planned to be released. Such a tool would be designed for researchers to quickly submit data that is expected to have some level of agreement and obtain their “*reproducibility score*”. This way protocols can be improved and reproducibility can be taken into account in the analysis.

Protocols for many bundles need to be developed for various purposes, such as clinical practice, synthesis of findings, building training sets for machine learning, etc. The segmentation plan and instructions need to be defined clearly by panels of experts, and agreed upon terminology [Mandonnet et al., 2018], to optimize reproducibility and anatomical validity. The field of manual tractography segmentation is decades behind fields such as grey nuclei or hippocampi manual segmentation on this matter. The latter has been refining segmentation protocols for the past decade and has already reached the state harmonized segmentation protocol and was evaluated with reproducibility in various settings [Boccardi et al., 2011, 2015; Frisoni et al., 2015; Apostolova et al., 2015; Wisse et al., 2017].

## 5. Conclusion

When trying to understand how similar WM bundles from dMRI tractography are, at least 3 values need to be taken into consideration: *Dice coefficient of voxels* showing how well the overall volume overlaps, *bundle adjacency of voxels* showing how far are voxels that do not overlap and *correlation of density map* showing if the streamlines are spatially distributed in a similar way. Results from our work on the pyramidal tract revealed that rater overlap is higher for voxel-wise measures (approximately 70%) than streamline-wise measures (approximately 35%). Distance between segmentations is lower for voxel-wise measures than streamline-wise measures, approximately 3mm and 6mm respectively. In comparison to the group average, the results depict an ease to identify true negatives, an adequate amount of true positives, while having a low amount of false positives. The voxel and streamline representations do not produce equal levels of reproducibility. Studies reporting bundle asymmetry in term of streamline count (streamline-based) will require a larger group difference than those reporting volume difference (voxel-based). This indicates a strong need for clear protocols for each bundle or at least detailed documents included with publications that used manual segmentation. Reproducibility of results is needed and goes hand-in-hand with the open science movement. A collaborative effort to evaluate and quantify human variability is needed.

## Acknowledgements

A special thanks to the funding sources for this work, the Fonds de recherche du Quebec – Nature et technologies (FRQNT) and Collaborative Research and Training Experience Program in Medical Image Analysis (CREATE-MIA) programs. Thank you to the Neuroinformatics Chair of the Sherbrooke University which helped push forward neurosciences research.

